# Astrocytes are active: An information theoretic approach reveals differences in Ca2+ signaling patterns among distinct astrocyte subtypes

**DOI:** 10.1101/2023.11.01.565176

**Authors:** Nicholas J. Mennona, Barbara Barile, Hoony Kang, Valentina Benfenati, Grazia P. Nicchia, Kate M. O’Neill, Wolfgang Losert

## Abstract

The discovery that astrocytes are an active, rather than a passive, component of the brain has ushered in a paradigm shift in thinking about how the brain processes information. Although the mechanisms by which astrocytes integrate information from neurons are still debated, such discourse should not distract from the importance of more completely understanding how astrocytes communicate via signals amongst themselves. This work aims to study how different astrocytes signal within their own networks. We investigate group calcium (Ca^2+^) dynamics in polygonal, stellate, and reactive astrocytes. These distinct and important astrocyte subtypes are present in the brain to varying degrees at different physiological states. We use an information-theoretic framework to quantify the dynamics embedded in the Ca2+ traces within astrocyte networks; specifically, we employ the Hurst exponent, cross-correlation, mutual information, and partitioned entropy to assess differences in the astrocyte signals across subtypes. To gain insights into the ability of astrocyte networks to respond to changes in the extracellular environment, we probe the networks with perturbations affecting their cytoskeletal dynamics (Latrunculin B) and energetic levels (Adenosine triphosphate). Overall, these three classes of astrocytes behave differently and respond idiosyncratically to their extracellular environment. We find that polygonal astrocytes are not quiescent, stellate astrocytes respond most strongly to ATP, and reactive astrocytes are uniquely perturbed by Latrunculin B. Interestingly, despite these distinct differences in behaviors, we find a uniform speed of information transport regardless of subtype or perturbation; this uniformity is maintained when using both cross-correlation and mutual information to assess this speed. We conclude that the differential ways astrocytes signal within our measured framework yield important insights into how astrocytes communicate and contribute to this pressing issue of understanding astrocyte information processing.

## Introduction

Astrocytes are the most numerous glial cell type in the brain. They are important for physiological functions, including K^+^ clearance^1^, glutamate homeostasis^2^, and extracellular volume regulation^3^ (reviewed in^4–6)^. Since astrocytes carry out these known supporting functions and lack electrical excitability, most studies have overlooked the potential active role of astrocytes in information processing. In contrast, neuronal dynamics has become synonymous with information processing^7–12^.

Nevertheless, recent work has provided strong indications that astrocytes do play an active role in information processing. As a counterpart to neurons, astrocytes also form their own networks, linked together via gap junctions composed of connexins, including Cx43, one of the main connexins in astrocytes^13,14^. Within these networks, astrocytes signal to each other also via slow Ca^2+^ waves.^15^. Intracellularly, IP_3_triggers intercellular Ca^2+^ waves^15^ which propagate through astrocyte networks via the aforementioned gap junctions. Recent studies reveal that these waves can occur spontaneously, with excitable dynamics similar to action potentials, although on a slower timescale of seconds^16^. Thus, astrocytes can have an active role in information flow, with calcium waves functioning as non-electrical communication signals. In single astrocytes, these calcium waves are heterogeneous^17^ and flow through astrocyte processes and microdomains^18–20^. Methods such as the machine learning tool AQUA now enable robust intracellular analysis of astrocyte calcium^17^ for *ex-vivo* and *in vivo* analysis of astrocyte activity.

Calcium signals, occurring either locally in microdomains or spreading into calcium waves also determine the release of bioactive molecules called gliotransmitters from astrocytes. Astrocyte gliotransmission (by e.g. ATP, Glutamate and GABA) is the known information transfer mechanism from astrocytes to neurons, and has been modeled for its role in synaptic plasticity, memory, and learning^21,26–28^. We propose that, to encompass such network scale information, astrocytes should also process and integrate information among themselves.

To support our hypothesis and gain insight on the information flowing between astrocytes and the unexplored network scale implications of that^16^, in this work, we utilize physics-based information-theoretic techniques to analyze and describe the behavior of calcium dynamics or pure primary rat culture of astrocytes *in vitro*. The current study contrasts three physiologically relevant subtypes of astrocytes, focusing on their collective (ensemble) dynamics within *in vitro* networks. These *in vitro* cultures remain a powerful tool^16^ for analyzing astrocyte dynamics, especially since the current study focuses on how individual astrocytes contribute to the ensemble.

Following established protocols, we study *in vitro* models of polygonal (immature), stellate (healthy), and reactive (immature) astrocytes. Intracellular and extracellular ionic changes are reported to occur upon several physiological and pathophysiological events, including regulation of the activity of chemical synapses, glial scarring^21^, synapsis plasticity^22^, and formation upon neurodevelopment^23^. Anisosmotic challenges are counterbalanced by water flows, which are predominantly mediated by the water channel aquaporin-4 (AQP4) in brain astrocytes^24^. AQP4 is reported to be polarized and enriched in astrocyte endfeet where it plays a key homeostatic role. By modulating water influx/efflux, it ensures a fine regulation of astrocyte cell volume and morphology, thus preserving brain integrity and functionality^24^. In this view, it is not surprising that AQP4 is found to be co-expressed or functionally coupled in astrocyte membranes with ion channels, including potassium (i.e. Kir4.1^25^) and calcium (i.e. TRPV4) channels^3,24,26^. On the other hand, connexins are shown to take part in astrocytes’ response to injury upon the stimulation of pro-inflammatory cytokines, including TGF-beta^27^. An increase in the levels of Cx43 in reactive astrocytes is proposed to act as a supporting mechanism that helps maintain astrocytes’ gliotic phenotype and maximize the coordinated intervention of these cells and their communication for the injury resolution^28,29^.

The astrocyte networks – polygonal, stellate, and reactive – are broadly classified as immature, healthy, and injured, respectively, and have distinct physiological behavioral profiles^19,30–33^. A key readout of these distinct subtypes is their unique and stark morphologies. The differing structures and morphologies of astrocytes, in general, have been implicated in brain functionality^30,34,35^ and the cell-cell signaling of astrocyte networks^36^. Notably, previous works evidence that different morphologies are paralleled by different functions in astrocytes, such as alteration in the cell volume mechanism^37–39^, electrophysiological properties linked with potassium spatial buffering^39,40^, and actin dynamics^41^. However, studies are lacking the capability of correlating spatially and temporally the calcium signaling in astrocytes, which are critically implicated in all the above-mentioned astrocytes functions.

With the goal of broadening our understanding of astrocyte-specific calcium signaling, we quantify the collective calcium dynamics, both spontaneous and perturbed, of these astrocytes. We perturb these dynamics *in vitro* cultures with low-dose Latrunculin B (LATB) and Adenosine triphosphate (ATP) as these represent basic perturbations to the cellular cytoskeleton^42–44^, and cellular energy^45–48^, respectively. Due to the slow and heterogeneous characteristics of astrocyte calcium signaling, astrocyte signals are more analog than digital. Neuronal signals are often treated as digital, 1 (firing) or 0 (not firing). Astrocyte signaling is too slow for such binarization. Thus, information-theoretic methods that extract information without digitizing the signal are used. We employ amplitude ordering symbolization with an embedding window large enough to encapsulate the dynamics of astrocyte calcium events, which are mostly on the order of 1-5.5 seconds^16^. Amplitude ordering, rather than binarizing via the mean, is more appropriate for quantifying the dynamic properties of these nonlinear signals. This nonlinearity allows us to elucidate information characteristics and information speeds across polygonal, stellate, and reactive astrocytes in a comprehensive manner.

## Methods

### Imaging

Primary astrocytes were obtained from Sprague Dawley rats housed at the University of Maryland (in concordance with the recommendations of and approval by the University of Maryland Institutional Animal Care and Use Committee; protocols R-FEB-21-04). Experiments were performed between 14 to 21 (DIV14-DIV21) days after dissection^49^. At this time, astrocytes were trypsinized from flasks and plated into PDL-coated (Sigma-Aldrich) 35 mm dishes (MatTek). Dishes containing cultured astrocytes were incubated with 5µM CalBryte 590 AM (AAT Bioquest) for 30-60 min at 37°C prior to imaging. Dishes were then washed with PBS. Astrocytes are plated at densities such that they proliferate to roughly 500k cells/well on the day of imaging (a final density of approximately 10k cells/mm^2^), and imaging was performed on a spinning disk confocal microscope (PerkinElmer). Image acquisition occurred for approximately 15 minutes at roughly one frame per second with 100 ms of exposure time per frame. The University of Maryland Imaging Core maintains the PerkinElmer spinning disk confocal microscope used for this research. The image sequences were taken by the equipped Hamamatsu ImagEM X2 EM-CCD camera (C9100-23B), which records 16-bit images. Sequences were taken from an air 20x objective (0.75 NA; 0.717 μm per pixel) and under temperature (37°C), CO2 (5%), and humidity control.

All experiments are performed in NeuroBasal media (Gibco). We use this media to simulate a neuronal environment, and we found that it allows for spontaneous activity in unperturbed wells (over DMEM). For simplicity, throughout this manuscript, this environment is referred to as spontaneous (control). ATP (Sigma-Aldrich) and Latrunculin B (Sigma-Aldrich) were applied immediately upon imaging and at low-dose concentrations of 1.5uM for ATP and 0.5uM for Latrunculin B. Unlike Latrunculin A, the slower depolymerization rate of Latrunculin B allows for perturbation of actin without drastically dissociating the entire astrocyte network. The value of Latrunculin was found based on testing whether our astrocyte networks were maintained (expressed minimal disassociation) for the 15-minute duration at the chosen imaging setting (e.g., laser intensity). For simplicity, throughout the rest of this work, we refer to Latrunculin B as LATB.

### Cell Culture

Primary astrocytes were obtained from Sprague Dawley rats housed at the University of Maryland (in concordance with the recommendations of and approval by the University of Maryland Institutional Animal Care and Use Committee; protocols R-FEB-21-04). Astrocytes were prepared from 0-to-2 day postnatal (P0-P2) Sprague Dawley rat brains as described previously^41^. Briefly, cortical tissue from each pup was separately dissociated via trituration, filtered, and plated into T25 flasks containing DMEM (ThermoFisher), 15% fetal bovine serum (Benchmark), and 1% penicillin-streptomycin (P/S). After roughly seven days, in order to generate stellate and reactive astrocytes, the media was changed to either (1) stellate media, which includes Neurobasal (ThermoFisher), 2% B27+ (GIBCO), 5 ng/mL HB-EGF (TOCRIS BioSciences)^19^ or (2) reactive media, which is serum-free DMEM to which 10 ug/mL of TGF-B1 was added after five additional days *in vitro*^50^.

### Cell lysis and Western Blot

Protein samples were obtained as previously described, with some slight changes^3^. Briefly, astrocytes grown for seven days in control, stellate, and reactive media were washed in ice-cold phosphate-buffered saline (PBS) and scraped with 300 µL of RIPA lysis buffer (10 mM Tris-HCl, pH 7.4, 140 mM NaCl, 1% Triton X-100, 1% Na deoxycholate, 0.1% SDS, 1mM Na3VO4, 1 mM NaF, 1 mM EDTA and 1× Protease Inhibitor Cocktail). Lysates were vortexed every 5 minutes for 30 minutes and centrifuged at 22 000 for 30 minutes at 4°C. Supernatants were collected and Pierce™ BCA Protein Assay Kit was used to dose the protein content. 5 µg of proteins/lane were dissolved in Laemmli Sample Buffer (Bio-Rad, Hercules, California, USA) added with 50 mM dithiothreitol, heated to 37°C for 10 min and resolved by SDS-PAGE in 10% (for Cx43 detection) and 12% (for AQP4 and GFAP detection) gels prepared with Mini-PROTEAN TGX Stain-Free Precast polyacrylamide solutions (Bio-Rad, Hercules, California, USA). gels were then activated for 5 minutes on a Chemidoc imaging system (Bio-Rad, Hercules, California, USA) and transferred to polyvinylidene fluoride (PVDF) membranes (Merck Millipore, Burlington, Massachusetts, USA). Stain-free signal of transferred blots was collected. Membranes were incubated with blocking solution and incubated overnight with primary antibodies, rinsed, then incubated with the appropriate peroxidase-conjugated IgG secondary antibodies. Protein-specific bands were revealed using enhanced chemiluminescent Clarity Western ECL Substrate (Bio-Rad, Hercules, California, USA) and visualized on a Chemidoc imaging system. The densitometric analysis was performed in Image Lab software 6.1.0 by normalizing the bands to the stain-free blot signal. The Western blot analysis of Cx43, AQP4, and GFAP for the polygonal, stellate, and reactive astrocytes confirms three distinct phenotypes: reactive upregulate Cx43 and AQP4, stellate upregulate AQP4, and polygonal cells do not express any of these markers significantly.

In this work, we investigate astrocytes of three physiological types; a prominent distinguishing characteristic of the differences is seen in the differences in morphology. Fig. 1 (top) provides representative images of the three classes of astrocytes analyzed in this study. Each astrocyte network does not contain a mix of different cell types and is cultured to maintain a singular physiological expression. For these astrocyte types, we investigate collective Ca2+ dynamics as illustrated in Fig 1 (middle), where multiple frames are overlaid to demonstrate the distribution of Ca^2+^ fluctuations (rising calcium in blue, diminishing calcium in yellow) in a network. Through Western blot analysis, we evaluated the expression profiles of 1) Cx43, one of the major isoforms forming astrocytic gap-junctions involved in calcium-mediated cell-to-cell communication, as a molecular marker of intercellular connectivity within astrocytes networks; 2) AQP4, the most abundantly expressed water channel in brain astrocytes, and GFAP, the predominant intermediate filament of the astrocyte cytoskeleton in the CNS, as proxy markers of cell differentiation and reactivity^3,51,52^.

**Fig 1:**
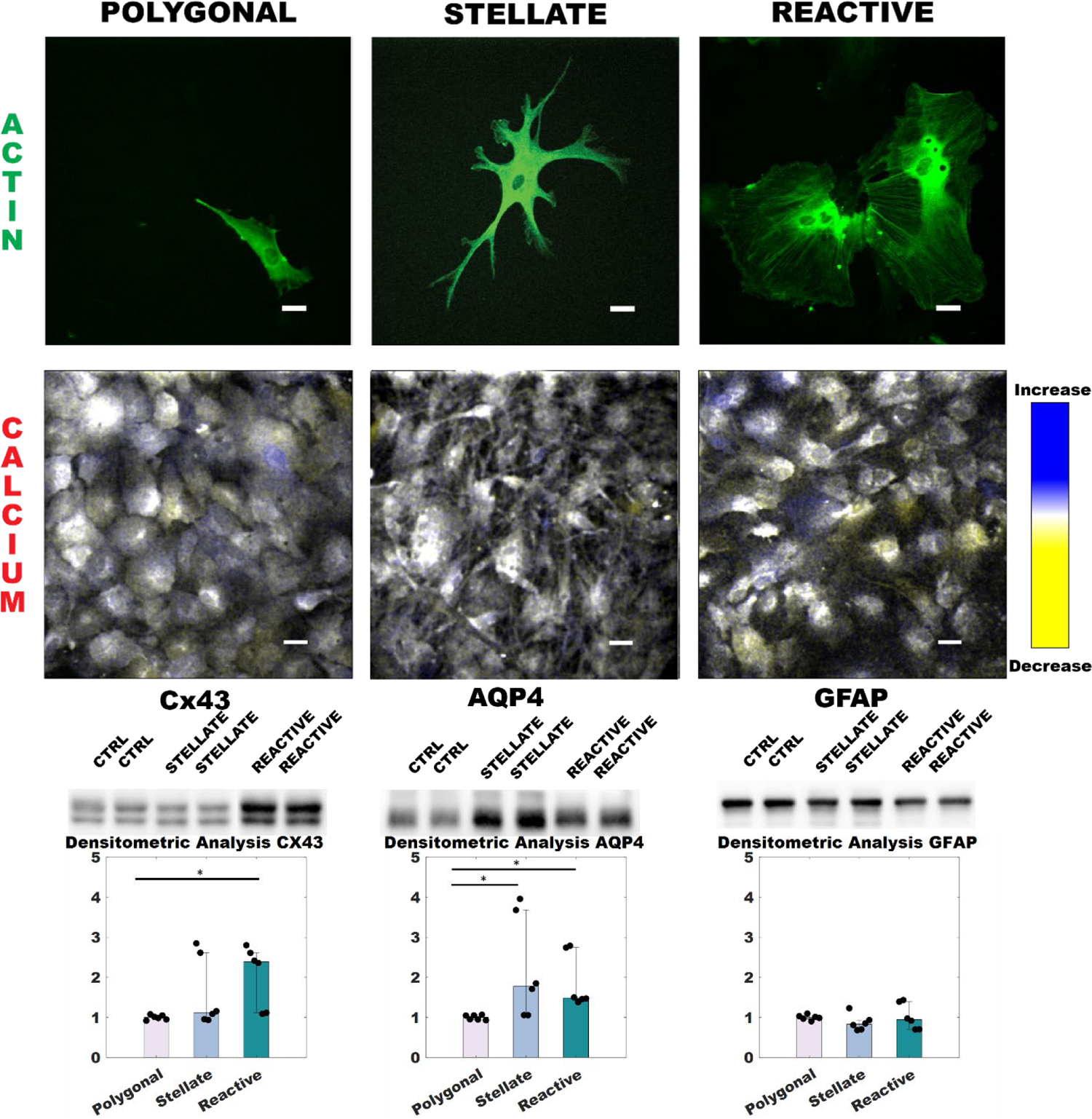
Astrocyte morphologies (polygonal, stellate, and reactive). The phenotypes correspond to different morphologies, as shown in green via actin images. Images of actin are acquired with 488nm light of astrocytes that express BacMam Actin-GFP. We record the calcium signaling channels of these dense astrocyte networks with 561 nm light of astrocyte networks stained with CalBryte 590. It has been noted that, regardless of protocol, different media formulations result in different morphologies for astrocytes^19^. We use time overlays to show the different manifestations of calcium propagation in the distinct morphological conditions of astrocytes. Blue corresponds to an increase in [Ca2^+^] during this frame, yellow a decrease, and white means that this frame contains values similar to the previous timepoint. Western Blot analysis of Cx43, AQP4, and GFAP in polygonal, stellate, and reactive astrocytes. The upper panels show Cx43 (band at ∼43kDa), AQP4 (band at ∼30-32 kDa), and GFAP expression (band at ∼50 kDa) in primary rat astrocytes. Western Blot analysis of Cx43, AQP4, and GFAP in polygonal, reactive, and stellate astrocytes. Western Blot showing Cx43 (band at ∼43 kDa), AQP4 (band at ∼30-32 kDa), and GFAP expression (band at ∼50 kDa) in rat astrocytes. Using Kruskal Wallis Dunn’s multiple comparison test, we find statistical significance for polygonal compared to stellate (AQP4) and reactive (Cx43 and AQP4). *p< 0.05; **p<0.01. Scale bars are 25 µm.

AQP4 and its supramolecular level of organization in larger-sized assemblies have been reported to correlate with the morphological and functional maturity of astrocytes in CNS postnatal development^53^ as well as in in-vitro models of astrocyte differentiation^3^. Therefore, the overexpression of AQP4 in stellate astrocytes is likely to refer to the morphological differentiation of this cell type in the experimental conditions. The overexpression of Cx43 in the reactive conditions might be ascribable to the acquisition in vitro of the in vivo-like reactive state that makes astrocytes more prompt at tissue healing and wound repair.

As for GFAP, besides the number of studies that identified a positive role for these canonical hallmarks of astrogliosis, just as many provided opposite evidence. On one hand, it was previously found to be downregulated in similar or even different models of astrogliosis (i.e., TNF and IL-1β^54–56^ and LPS-treated astrocytes^57^. On the other hand, its upregulation was associated with beneficial and neuroprotective effects i.e., enhancing neuronal regeneration through protein re-localization, increasing the membrane retention of glutamate transport GLAST in astrocytes, which protects surrounding neurons from glutamate excitotoxicity^58^; providing scaffolds for new neurites^59^. In the present study, no changes were detected in the three conditions, although we could detect differences in other markers in response to inflammatory (i.e., Cx43) and differentiation stimuli (i.e., AQP4; Cx43). We, therefore, anticipate that GFAP might not be a reliable marker to assess astrocytes’ degree of astrogliosis or differentiation.

Western blot analysis shows that the three cell types have distinct expression profiles of the aforementioned astrocyte proteins. Of note, we found that Cx43 is upregulated in the reactive state compared to control and stellate astrocytes (Fig.1, bottom left panel), while AQP4 is overexpressed both upon stimulation with TGF-beta and in star-shaped astrocytes, differently from controls (Fig.2, bottom middle panel). GFAP was equally abundant in the three conditions with no statistical significance (Fig.1, bottom right panel).

**Fig 2:**
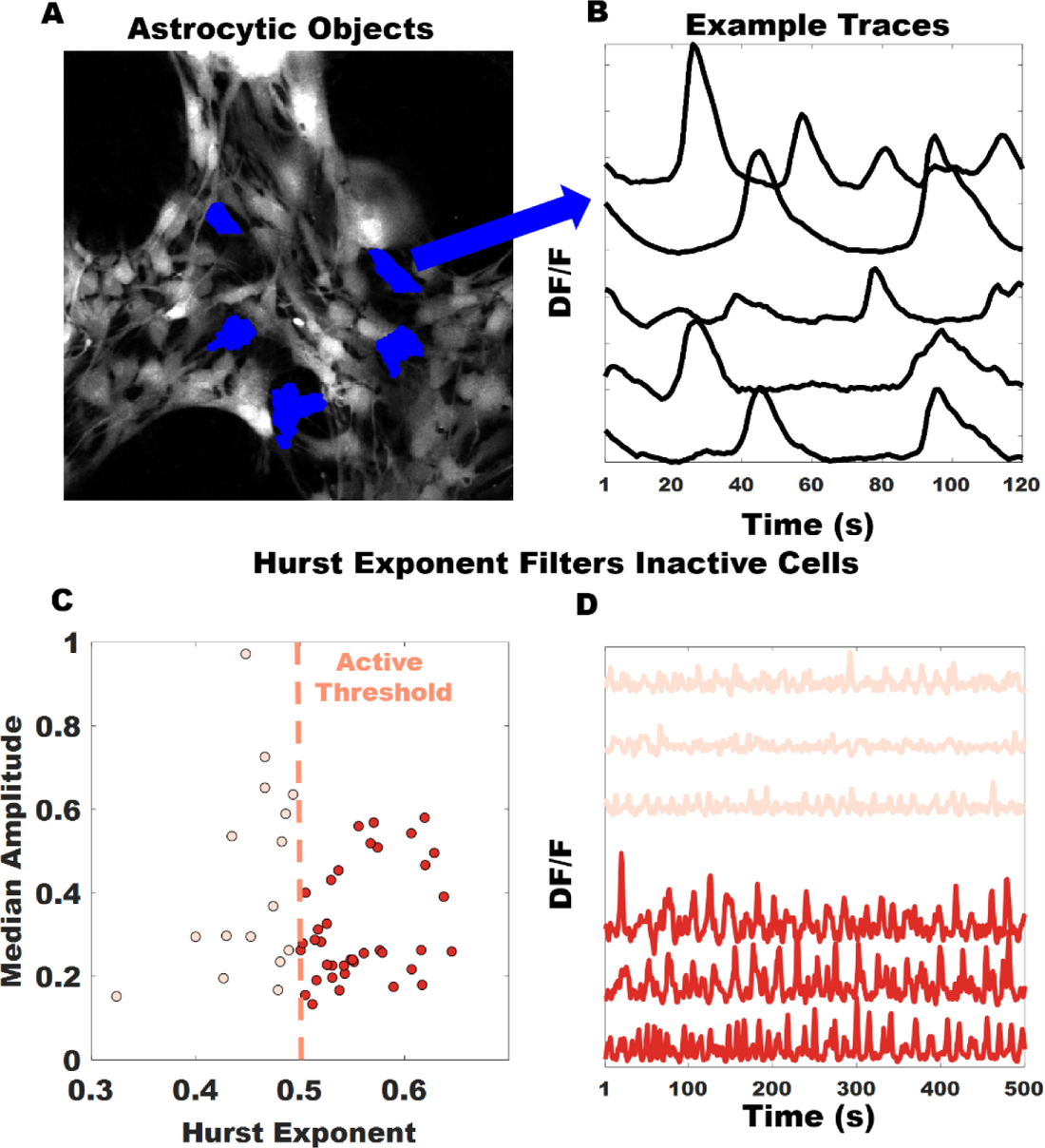
Astrocytes traces extracted and filtered using Hurst exponent. (A) Given a dense astrocytic network, astrocytic elements are segmented using FogBank. Examples of astrocytic elements extracted. (B) DF/F traces of the five objects extracted in blue (dark pixelated objects) are shown. (C) Hurst exponent measures memory in a time series, which is used to filter out inactive cells or poorly segmented objects. Inactive (salmon, light) versus active (red, dark) cells plotted. Only time series with Hurst exponents greater than or equal to 0.5 are used. (D) From the example points, we plot representative inactive and active traces.

Overall, while illustrating that the proteins undergo distinct changes in their profile expression in response to the different external stimuli provided under culture conditions, these results also provide evidence that the three cell types can be regarded as distinct “phenotypes” from the molecular perspective.

### Analytical Methods

To analyze the calcium dynamics of astrocytes, we first segment individual astrocyte objects within our imaging datasets using the morphological watershed algorithm FogBank^60^. Manually, we refined parameters per each imaging sequence such that segmented boundaries roughly correspond to the distinctions made by eye. FogBank preserves the unique heterogeneity of astrocyte bodies rather than using the cell center with an arbitrary radius, as is typically done with neuron-imaging analysis^61^. See Supplement for a representative FogBank example. All image sequences are jitter corrected, and Gaussian smoothed in the spatial domain before further analysis. For each frame in a dataset, we take the median fluorescence value from all the pixels identified within the astrocyte object, as shown in the five objects in Fig 2A. We compute relative fluorescence measure (ΔF/F_0_), where ΔF=F-F_0_, for each astrocyte object. F_0_ is computed using a window of approximately 30s (30 frames, 15 back, 15 forward) where F_0_ is the average of the fluorescence values below the 50^th^ percentile of values contained within the window^62^.

This is called a trace. Throughout the paper, for convenience, we address this fluorescence measure as DF/F. As seen in representative traces, peaks in calcium dynamics are broad and not well-defined. The sliding window is chosen based on an inspection of the data and does not meaningfully change the dynamic properties of the traces. Please consult the Supplement for more details. We chose this window size based on experimental observation of astrocyte calcium event duration. Examples of such traces are shown in Fig 2B. Recall that astrocytes are not neurons, so it is inappropriate to call the activity of our traces ‘spikes.’ Instead, we use the term “calcium event” to describe the smooth rises and falls exhibited by traces in Fig 2B.

Our goal is to use traces that show persistent calcium signals and to distinguish them from random fluctuations of signals in the networks. Our goal is to define traces that exhibit smooth and meaningful rises and falls (see red, darker curves in Fig 2D) as ‘active,’ regardless of the intensity values. The Hurst Exponent (defined as *H*) allows us to carry out this analysis. H is defined for a given time series x(t) as:

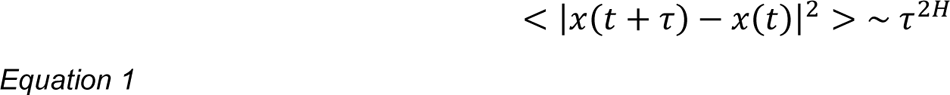

We find that the Hurst exponent provides a robust and reliable way of filtering out noisy traces due to poor segmentation or weak calcium signals. Hurst exponent values below a threshold of 0.5 represent random activity, and the corresponding traces (e.g., the salmon, lighter colored curves in Fig 2D) are excluded from the analysis^63,64^.

The importance of correlations in neuronal encoding is well-cited in the literature^65–68^. The study of correlation in the brain relates to the collective activity of neuronal ensembles in relation to information encoding and processing in the brain. This level of mathematical rigor has not been applied to astrocyte-only networks, and the study of correlation within astrocyte networks has only been viewed in the context of neuronal activation^69^. Given the success of using these methods on neurons and the absence of their use in analyzing astrocytes, we argue that our use of these methods should yield comparable success and insight. Correlations allow us to understand the global characteristics of an ensemble of active nodes (in this case, astrocytes). However, we can alternatively view the dynamics in our astrocyte networks as a dynamical system that is affected by individual astrocytes, i.e., the local dynamics of one trace affecting the global dynamics observed. Since our Western blot analysis confirms cell-cell coupling, we argue that these readouts provide a fairly accurate representation of the information processing performed by these collections of astrocytes. This approach has been well-documented experimentally (e.g., with neuronal MRI data) and theoretically^70–78^. A limitation of correlations is that the global characteristics of a network are studied by assessing each cell’s connections to others in the network. Information theory helps overcome such limitations. An information theoretic approach can analyze a network for each astrocyte rather than for pairs of cells. Thus, this approach provides a robust metric for assessing each individual astrocyte’s contribution to the mean field or the group characteristics of the network.

To implement this approach, from the extracted traces, we convert the traces into states using symbolization^70,79^. This symbolization enables the interpretation of changes in calcium as an analog signal; traces that exhibit lower entropy correspond to traces with more periodic fluctuations. We take the time traces shown in Fig 3A, the amplitude order of each timepoint in a window, and convert this set of timepoints into a state (an example suite of states is available in Fig 3B), and then convert the traces into symbol sequences visualized in Fig 3C. As an illustrative example with a manageable number of states, m=3 has been used (number of states = m!). From the resulting symbol sequence, we can find the probability distributions of states (Fig 3D). The probability of states, in turn, allows us to compute an information entropy for this representation of calcium traces. Amplitude ordering entails choosing two parameters, m, and l, where m defines the embedding dimension^70,78^ (the symbol set size that functions as a coarse-graining window) and l denotes the time lag^78^. A more comprehensive review of this symbolization method can be found in Refs^70–72,74,76–79^. We choose m to smooth out noise without losing actual spikes in the data. Based on our experimental observation of the duration of calcium events in our data, we chose m=5 and l=1. As m takes the role of the coarse-graining window, l can be kept at its minimum (l=1) to prevent further information down-sampling.

Given that we have converted these time series into symbol series, we can perform time-delayed mutual information methods. Time-delayed mutual information is the symbolized equivalent of cross-correlation. Moreover, given that astrocyte signals are nonlinear, mutual information is a more appropriate method than Granger causality^80^. While there are some similarities between mutual information and cross-correlation, we emphasize that mutual information discusses the dynamics of symbols (with embedded dynamics) from a probability distribution. Cross-correlation looks at the absolute fluctuations from the mean for traces. In some regards, then, symbolization can be viewed as the differential expression of trace dynamics. Lastly, from both of these measures, we can quantify the speed of information transport^81^. We can measure speed from the measured distances between astrocytes and either the cross-correlation time lag or the time lag associated with the mutual information measurement.

Additionally, we analyze the information content of astrocyte networks with partitioned entropy. Partitioned entropy h_a_(τ) is a useful metric for understanding the time-evolution of states over a certain

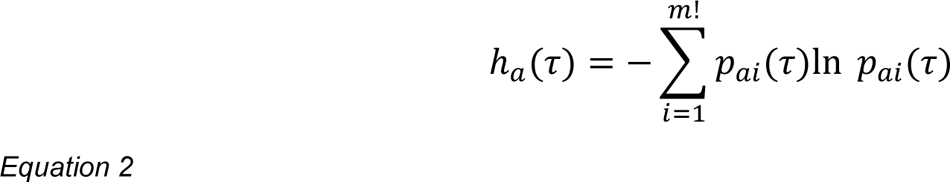

where p_ai_ (τ) represents the probability that the system evolves from state a to state i at τ timesteps. Partitioned entropy differs from canonical entropy as it is a function of time. We employ this approach to understand better how the ramps and falls of these calcium fluctuations evolve differentially for all of the cells within the unique networks. A schematic for understanding partitioned entropy is shown in Fig 3E. We validated this analysis on both regular and chaotic time series data.^72^ See Supplement for details. As a clarification, we note that the symbolization described enables analysis of Shannon entropy. The rest of the analysis refers to Shannon entropy and the dynamics therein, e.g., partitioned (Shannon) entropy.

**Fig. 3:**
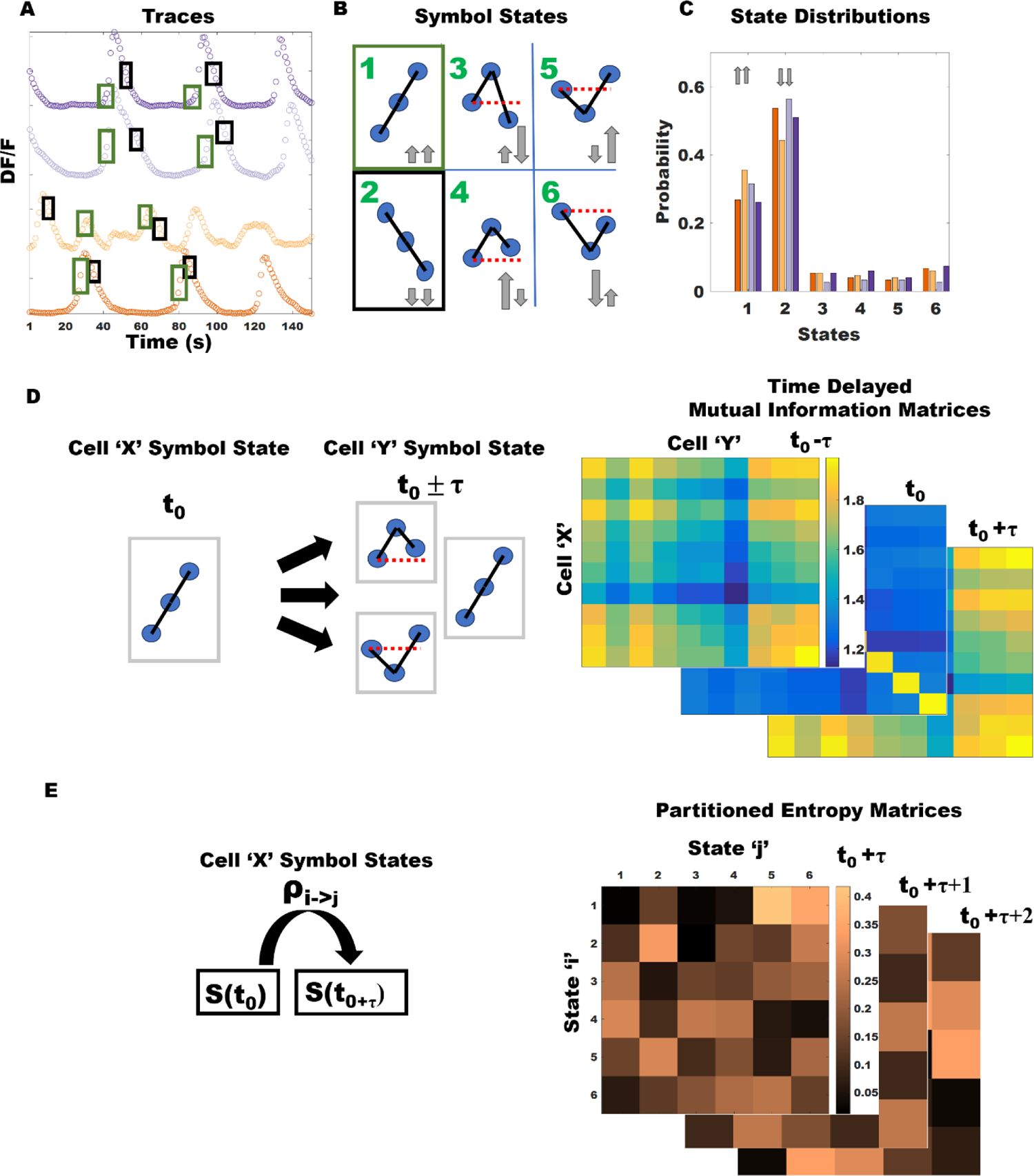
Methods for Information Theoretic Approach to Collective Astrocyte Dynamics. (A) Astrocyte traces. Different shaded boxes correspond to example symbol states. (B) Symbol states. Shaded boxes are those previously overlaid on traces. In this figure, have used an m of 3 as this corresponds to 6 possible states for every time window (m!=6). We note, using grey arrows, the physical dynamics behind each state (for states 1 and 2, the same size for the arrows is for visual purposes). (C) Symbol sequences for the traces shown previously. (D) Visualization of Time-Delayed Mutual Information. (E) Example subset of transition probabilities from states to future states for some given tau for any particular state ‘i’. See Equation 1 for more details. This subfigure represents partitioned entropy.

## Results

### Astrocyte Speed of Information Transport

An important metric for assessing how astrocyte transport information is information speed (speed of information transport). This ad-hoc (average) metric is extracted for free by any time lag calculation. We do not track calcium flow, but can track the exchange of information as a function of time and distance between cells. This method has been used for mutual information^81^; we extend it for cross correlation since the time lag associated with maximal cross correlation is similarly extracted. Our measure of speed is average (*v= DIST/τ),* it coarsely reveals the rate at which these pairwise quantities are absolutely maximized, reflecting the nature of the timescales associated with astrocyte calcium events. Additionally, the maximal time lags found are rate-limited by the acquisition speed, which is 1 Hz. Due to the 1 Hz frame rate and other coarse graining involved (e.g. distance is computed from center of mass points in a segmented image), this measure is an average metric. Thus, we conclude that a few micron per second difference is not significant to ascertain any meaningful discrepancy from a biological perspective.

We describe the cross-correlation speeds first. The distributions are modelled as a lognormal distribution to obtain the mean speeds to follow as shown in Figs 4A-C. Outliers at infinity are removed (which may happen for time lags of zero). We report the mean of these distributions modeled under lognormal assumptions. The spontaneous speeds are: for polygonal 25 µm/s, for stellate 28 µm/s, for reactive 23 µm/s. Under LATB, for polygonal 25 µm/s, for stellate 27 µm/s, for reactive 21 µm/s. With ATP, for polygonal, 29 µm/s, for stellate 28 µm/s, for reactive 25 µm/s. As expressed in Figs 4A-C, these distributions are not symmetric (not entirely lognormal). Given that the order of difference between stellate and other networks is ∼5 µm/s at most, and the distributions themselves retain similar characteristics (the distributions peak within this 20-30 µm/s range consistent with experimental studies), we conclude that the speed of information transport is invariant across astrocyte subtypes and conditions.

**Fig 4:**
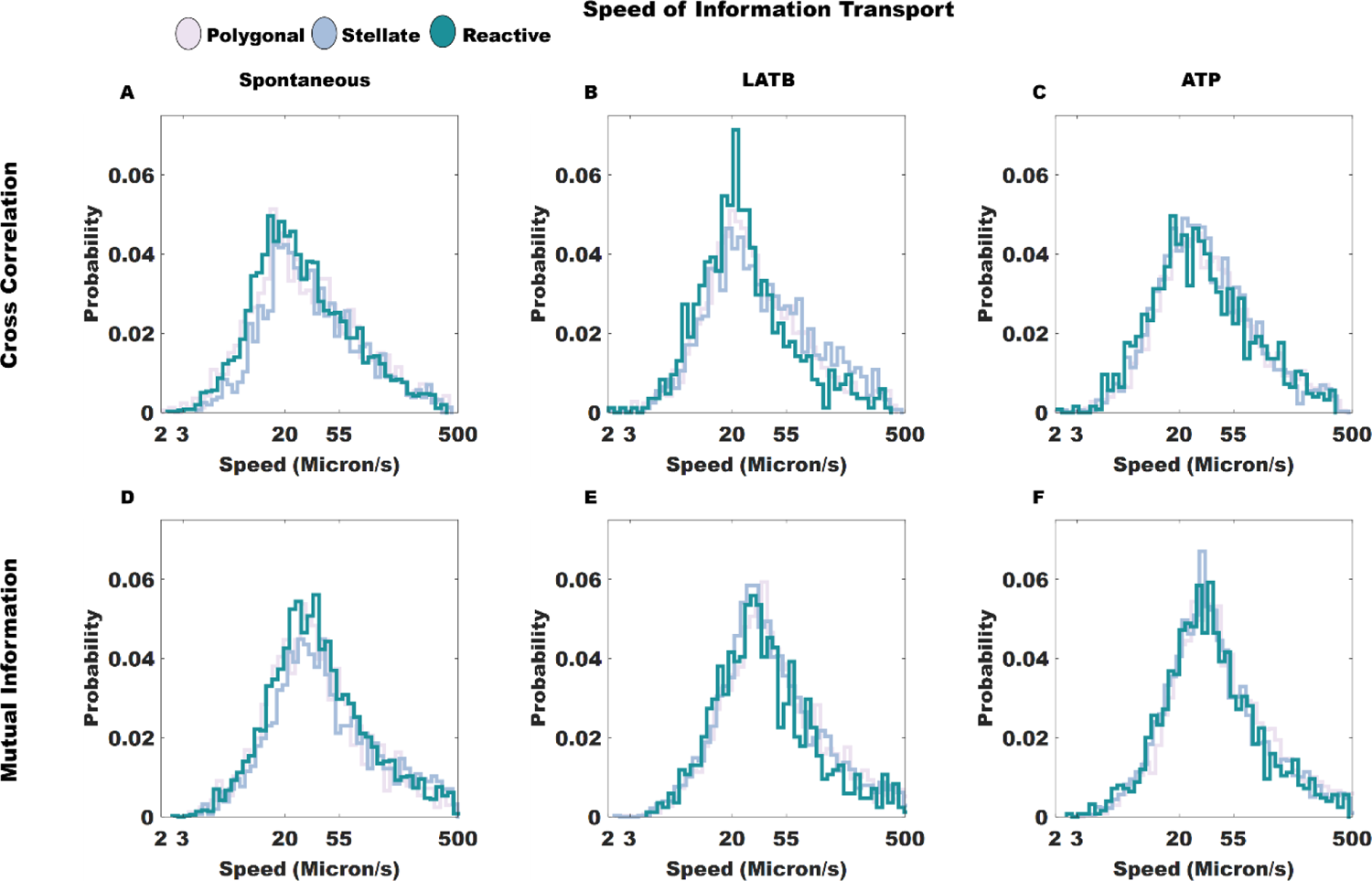
Speeds of Information Transport via Cross Correlation and Mutual Information. Figures shown for all distinct subtypes of astrocyte. Cross correlation speeds are shown for (A) spontaneous, (B) ATP, and (C) LATB affected networks. Mutual information speeds are displayed for (D) spontaneous, (E) ATP, and (F) LATB affected networks. Due to the skewed nature of all distributions, these distributions have been log-normalized. We find the speed of information transport is invariant across astrocyte subtype and extracellular environment.

Next, we report the mutual information speeds. The spontaneous speeds are: for polygonal 28 µm/s, for stellate 32 µm/s, for reactive 30 µm/s. Under LATB, for polygonal 34 µm/s, for stellate 32 µm/s, for reactive 29 µm/s. With ATP, for polygonal 34 µm/s, stellate 30 µm/s, and reactive 29 µm/s. These mean values are reported from modelling the distributions as lognormal. We conclude that the apparent differences between the reported mean values are minimized for mutual information speeds. The speeds are slightly higher than those of cross correlation. All speeds are in the ∼30-35 µm/s. This range is roughly consistent with the cross-correlation speeds (at the higher end), and is consistent with the reported literature.

Largely, we find a constant speed of information transport across conditions and types. Using this metric (defined as distance between objects divided by the time lag associated with maximization of pairwise quantity), we find that all distributions sharply peak similarly within the reported experimental range for calcium wave speeds. The nature of these distributions across both subtype and conditions suggests an invariance of the speed of information transport in astrocyte networks using this coarse approximation. The similarities in the relationships between the distributions for the quantities (mutual information and cross-correlation are similar, but differently formulated) reflects that the aggregate relationships between calcium signals for astrocytes are independent of subtype and perturbation. This counter-intuitive result suggests that astrocytes regulate ions’ flow despite cellular state, and maintain homeostasis in stressful environments to maintain signaling speeds.

As a final remark, we comment that the time-lags associated with either maximal cross-correlation or mutual information may come from a symmetric peaking in a time-lag curve (i.e. bimodal peaks about the y-axis). We argue that many of the time lags reflect a maximal value over a relevant physiological window. All other lags reflect a mode within the time-lag curves themselves. This argument is expanded in the Supplement.

### Information exchange of signaling amongst active astrocytes

Next, we assess how the traces in astrocyte networks can be used to infer relationships across pairs of cells. Given a time window of 31 frames (roughly 31 s), we find the maximum cross correlation and mutual information values. These relationships are shown in Figs 5A-C. First, we discuss the cross-correlation distributions. In spontaneous conditions, polygonal astrocyte pairs have higher cross correlation subtype, but stellate and reactive pairs demonstrate comparable distributions (Fig 5A). When LATB is introduced into the astrocyte networks; the distributions are separable indicating that the breakdown of the actin cytoskeleton causes idiosyncratic (i.e. each astrocyte network varies its distribution relative to its spontaneous profile) changes to astrocyte subtypes (Fig 5B). When ATP is introduced, all distributions curves aggregate (Fig 5C). The effect of ATP suggests a minimization between the global differences in astrocyte populations. This minimization is reflected in the strongly overlapping distributions.

**Fig 5:**
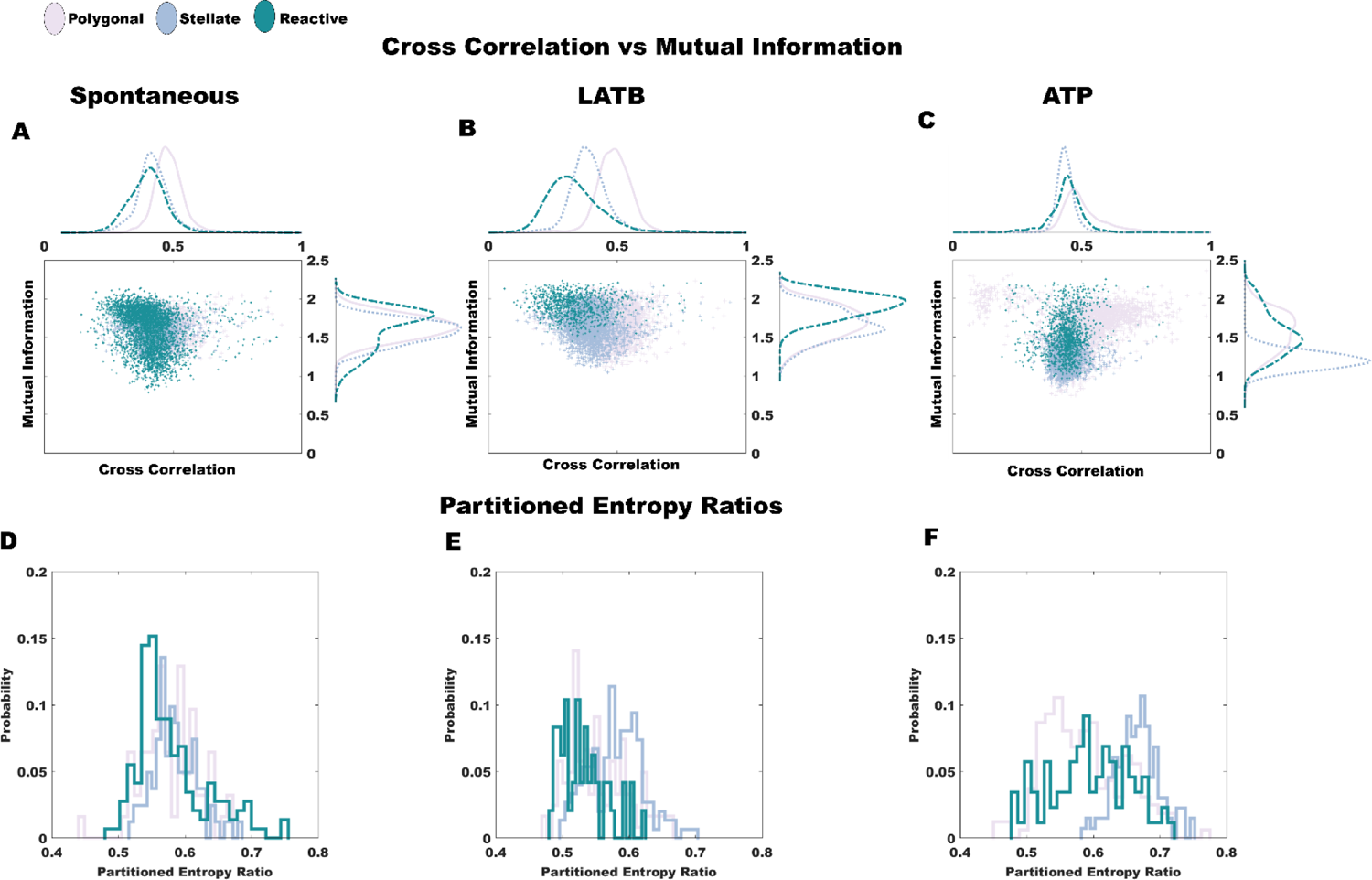
Cross correlation versus mutual information, and partitioned entropy comparisons. We compare the distributions of both maximal cross correlation and maximal mutual information across time for (A) spontaneous, (B) LATB-induced, and (C) ATP-enhanced astrocyte networks. Each point represents the cross correlation (x-axis) and mutual information (y-axis) for the same pair of cells. We then look at the dynamics of individual traces using partitioned entropy for (D) spontaneous, (E) LATB-induced, and (F) ATP-enhanced networks. Partitioned entropy ratio is the average of the partitioned entropy divided by the entropy.

Next, we discuss mutual information. In spontaneous conditions, we find that the distributions for the astrocyte polygonal and stellate astrocytes are similar (and lower than reactive). These results suggests that spontaneously polygonal and stellate astrocyte behave more independently than reactive astrocytes (Fig 5A). When LATB is introduced, all mutual information curves are shifted higher with reactive astrocytes retaining larger values while polygonal and stellate astrocyte pairs are similar (Fig 5B). Interestingly, this result indicates that a breakdown in the cytoskeleton causes uniformity in local behavior in astrocyte interactions (i.e. higher values of mutual information reflect less independence). When ATP is introduced, stellate astrocytes respond the most strongly (Fig 5C). Stellate astrocytes become strongly independent in terms of their local behavior. While stellate astrocytes respond the most strongly, all astrocyte subtypes become more independent (lower mutual information) when exposed to ATP.

### Partitioned entropy highlights temporally local dynamics

We look at the partition entropy ratios for astrocytes in networks. Unlike with cross correlation and mutual information, partitioned entropy is a method for analyzing the dynamics of individual traces themselves (i.e. it is not a pairwise analysis method). As a dynamical measure, it provides more ability for understanding information dynamics than reporting entropy values alone. Partitioned entropy ratios are calculated by finding the average partitioned entropy across some time window (in this paper we chose to start at τ=10 and end at τ=150, see Supplement for details). Values of partitioned entropy closer to 1 are classified as chaotic whereas values closer to 0.5 are deemed limit cycles (e.g. sinusoidal oscillations). Classifying astrocytes in this way is beyond the scope of this work, but it is useful for further distinguishing dynamical profiles of the subtypes.

We find that spontaneously, astrocytes retain similar individual characteristics, as demonstrated in Fig 5D. This result is surprising given the numerous reported physiological differences in these subtypes. Coupled with some similarities across subtypes in the pairwise interactions results, we argue that, in spontaneous conditions, when astrocytes signal it is similar. This result suggests a universal characteristic to spontaneous astrocyte calcium events. LATB, comparable to the distinct effects in pairwise interactions, causes reactive astrocyte to behave more sinusoidally (Fig 5E). Stellate astrocytes respond with higher partitioned entropy ratios. Lastly, where LATB induces the strongest separation in cross-correlation distributions, ATP seems to induce a comparable outsized effect in the partitioned entropy curves shown in Fig 5F. The different perspective (focusing solely on the individual embedded dynamics) provided by partitioned entropy complements the results mentioned above in ascertaining the nuanced differences in astrocyte calcium signaling.

### Wasserstein distances enhance interpretability of comparisons

We use the Wasserstein distance (using its L1 norm definition, Wasserstein-1) to measure the differences in distributions shown in Fig 5. Wasserstein-1 quantifies the minimum distance needed to shift all points in a distribution to match another distribution. The Wasserstein distance considers the individual values used to generate the histograms in Fig 5., and is independent of the choice of bin. We can measure a Wasserstein distance not only to compare distributions, but also to characterize a single distribution by using bootstrapping. Bootstrapping was performed for all Wasserstein-1 calculations with a sample size of 100 and 10,000 iterations. We can calculate Wasserstein distances for all metrics, cross correlation, mutual information, and partitioned entropy with this approach.

Under spontaneous conditions, polygonal cross correlation values are the most distinct (Fig 6A). Using symbolization, we find that the differences between stellate and reactive astrocytes (S-R) become most apparent for mutual information (Fig 6B) and partitioned entropy (Fig 6C). Thus, we conclude that on a global interaction perspective, polygonal dynamics behave the most uniquely; on a local level (using the embedded symbol states to understand dynamics), we find that spontaneously stellate astrocytes have a unique dynamical profile relative to the reactive subtype. The ability of information theory to tease out this result is significant. Both stellate and reactive astrocytes possess branches^82^. The process formation could have implicated such processes in the similarities seen in their cross-correlation distributions (S-R of Fig 6A).

**Fig 6:**
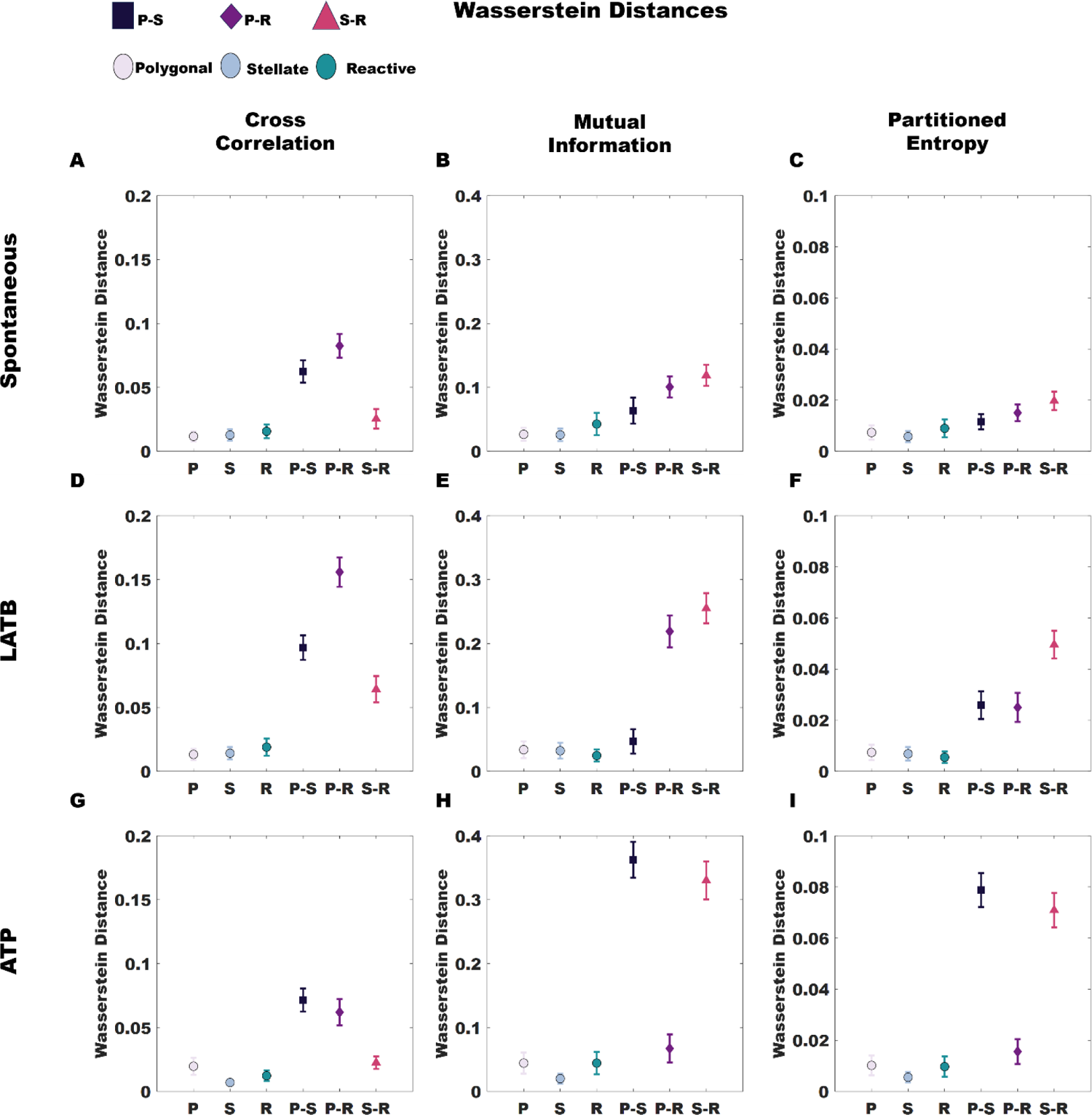
Wasserstein distances for the three classes of distributions shown in Fig 5. Astrocytes types are grouped; three pairs exist as highlighted by the dark (polygonal-stellate), purple (polygonal-reactive), and pink (stellate-reactive). We contrast these groups against the natural variance within the individual subtype distributions. We find the Wasserstein distances for: spontaneous (A) cross correlation (B) mutual information, and (C) partitioned entropy distributions, LATB exposed (D) cross correlation (E) mutual information, and (F) partitioned entropy distributions, and ATP induced (G) cross correlation (H) mutual information, and (I) partitioned entropy distribution. Results are bootstrapped.

Relative to spontaneous conditions, LATB increases the differences in the distributions for cross-correlation (Fig 6D), mutual information (Fig 6E), and partitioned entropy (Fig 6F). Where again, we see that polygonal astrocytes on a global level are the most distinct relative to stellate and reactive astrocytes (Fig 6D), but we find that in the mutual information distributions, P-R and S-R, reactive astrocytes separate the most strongly. This suggests reactive astrocytes are more susceptible to cytoskeletal perturbation, limiting pairwise signal propagation. Importantly, the high Wasserstein distances in Figs. 6D and 6E distinguish polygonal and reactive (P-R) astrocytes despite the similarities in AQP4 expression. The strongest separation in terms of individual trace dynamics is seen in S-R (Fig 6F), suggesting a strong difference in effect when healthy (stellate) versus unhealthy (reactive) astrocytes are exposed to LATB.

Lastly, it is interesting that the highest reported Wasserstein distances for mutual information (Fig 6H) and partitioned entropy (Fig 6I) are found under exposure to ATP. The lowest differences are attributed to the cross-correlation (Fig 6G). Outside of physiological comparisons, this highlights the importance of the diverse set of information theoretic tools used for understanding astrocyte signal dynamics. In contrast to the effect of LATB, ATP appears to cause the entire network to become excited (i.e. hyper-active), consistent with its known role as a key gliotransmitter. The lowest Wasserstein differences (across all conditions) are reported for P-R. Indeed, this result indicates that stellate astrocytes are the most susceptible to ATP intervention (P-S and S-R in Figs. 6H and 6G) whereas polygonal astrocytes share the unhealthy characteristics of reactive astrocytes (P-R).

## Discussion

Collective astrocyte dynamics are rich in information distinct from neurons and previously overlooked. Starting from filtering out inactive traces with the non-biased Hurst exponent, we employed cross correlation, mutual information, and partitioned entropy to ascertain the dynamical properties of collections of maturing (polygonal), healthy (stellate), and injured (reactive) astrocytes. This behavior characterization extends to these astrocyte networks defined by distinct states (subtypes defined by differences e.g., in molecular expression, morphology) and when gliotransmitters or biomechanics (LATB) of the system is altered. Interestingly, this focus on information differences allowed us to demonstrate uniformity in the speed of information transport across astrocyte subtype and extracellular environment (ATP/LATB). Our analysis on these astrocyte networks handles the complex spatiotemporal patterns present in these data. Not merely regarding the calcium signaling of astrocytes as an adaptive mechanism, our work argues that the spatiotemporal patterns embedded in these networks are active. That these patterns themselves are sources of dense information rivaling neuronal networks. The spatiotemporal patterning of spontaneous astrocytes is shown in the Supplement.

A principal finding of this work is the corroboration of astrocyte signaling (Ca2+ waves) speed using information theoretic methods. Using cross correlation and mutual information on astrocyte traces, aggregating traces from independent image sequences, we find an invariance of the speed of information transport. We discovered that for all conditions investigated, the speed of information transport is approximately 25-40 µm/sec, consistent with the speed of Ca^2+^ waves measured directly in prior studies ^49,83,84^. These findings are independent of astrocyte subtype and chemical milieu, which strongly indicates that Ca^2+^ waves are the primary mechanism of astrocyte-to-astrocyte information exchange, i.e. Ca^2+^ dynamics is a dominant form of communication in astrocyte networks for all states and environmental conditions investigated despite differences in molecular expression, information processing, and the biochemical cell-cell coupling machinery (AQP4, Cx43). Given that we will subsequently discuss the differences in information dynamics, this result is even more startingly. Confirmation of the speed of information transport with previously reported values lends confidence to using these methods for understanding astrocyte signals.

We preface subsequent interpretations of our results with subtle distinctions between the competing pairwise information theoretic concepts in this work. Correlation (of which cross is the most appropriate for astrocyte signals; see Supplement for more details) deals with the specific similarities of mean field fluctuations of the traces. One way to understand this subtle concept is that pairwise correlation values relate to the absolute coarse-grained differences between points in a time series. Accounting for the time lag to maximize correlation, in the context of astrocyte signals, it describes how similar the calcium events look; the average area overlap of all calcium events. Mutual information, supported by symbolization, is more focused and able to provide more local descriptions of how symbol state influences across time series^85^Both are important, and our results, such as the marked separation between cross-correlation LATB distributions, indicate that both provide a holistic ability to compare astrocyte dynamics. Indeed, an externality of these results, including the use of partitioned entropy for comparative analysis, is a demonstration of the varied insights that are gained by using this complete set of information theoretic tools.

Importantly, polygonal astrocytes in control conditions (spontaneous), which have been regarded as quiescent,^19^ exhibit information processing; in other words, polygonal cells are not inactive cells and have complex signaling patterns. The difference between these polygonal cells, and stellate and reactive astrocytes at first lies in the global correlation structure amongst polygonal astrocytes. As shown in Figs 6A,6D,6G, polygonal cells are the most distinct with higher cross correlation values than stellate and reactive astrocytes. As correlation is a global averaging of time values, this suggests that the mean field behavior of polygonal cells is more coupled than those of stellate and reactive astrocytes. In other words, polygonal astrocytes broadly have more global network synchrony. These insights are counterintuitive as the higher expression of the astrocyte markers regulating cell-cell coupling, AQP4 and Cx43, seen in stellate and reactive astrocytes (Fig 1) suggest that these cells would couple more closely in their information processing. One possible explanation is that the lower expression of coupling proteins limits the signal propagation to fewer polygonal neighbors and thus, paradoxically, these polygonal networks are globally more aligned. Future studies will investigate this possibility.

Locally, we gain additional insights into stellate and reactive signaling dynamics using symbolization which embeds the dynamics. We conclude that reactive astrocytes respond most strongly to cytoskeletal perturbation (LATB) whereas stellate astrocytes react most strongly to gliotransmitter ATP. These results are consistent with the role of the cytoskeleton and ATP in reactive and stellate astrocytes, respectively. Reactive astrocytes must have a dynamic cytoskeleton as they migrate to form the glial scar, an adaptive function in response to injury or disease^35,86^. Stellate astrocytes communicate via more localized contacts, and ATP is suggested to be an important extracellular messenger for specifically stellate-like calcium waves^87,88^.

Overall, our work shows that applying information theoretic tools to *in vitro* astrocyte networks lends insights into the collective action of astrocytes. Generally, our work has analyzed the complex spatiotemporal characteristics of polygonal, stellate, and reactive networks. We find that polygonal astrocytes are not quiescent, stellate astrocytes respond most significantly to ATP, and reactive astrocytes are most affected by LATB. Our study also provides a systematic baseline of three complementary astrocyte states and key perturbations supporting future studies of astrocyte network function in health and disease. The successful demonstration of our method across numerous astrocyte conditions and microenvironments suggests broad applicability and promise for these tools to study astrocyte dynamics.

## Acknowledgments

The authors thank the University of Maryland Imaging Incubator Core Facility for providing and maintaining the systems used in collecting images for this work. This work was supported by US Air Force Office of Scientific Research Astro-COLL grant FA9550-21-1-0352. We want to thank Dr Leonard Campanello for always being generous with his time and providing insightful guidance on the timescales of information. We additionally thank Prof Pratyush Tiwary for his comments and suggestions.

## Software Availability

Software has been made available on GitHub at: https://github.com/losertlab/Astrocyte_CollectiveNetworkInformation

