## Supplement for "Astrocytes are active: An information theoretic approach reveals differences in Ca2+ signaling patterns among distinct astrocyte subtypes"

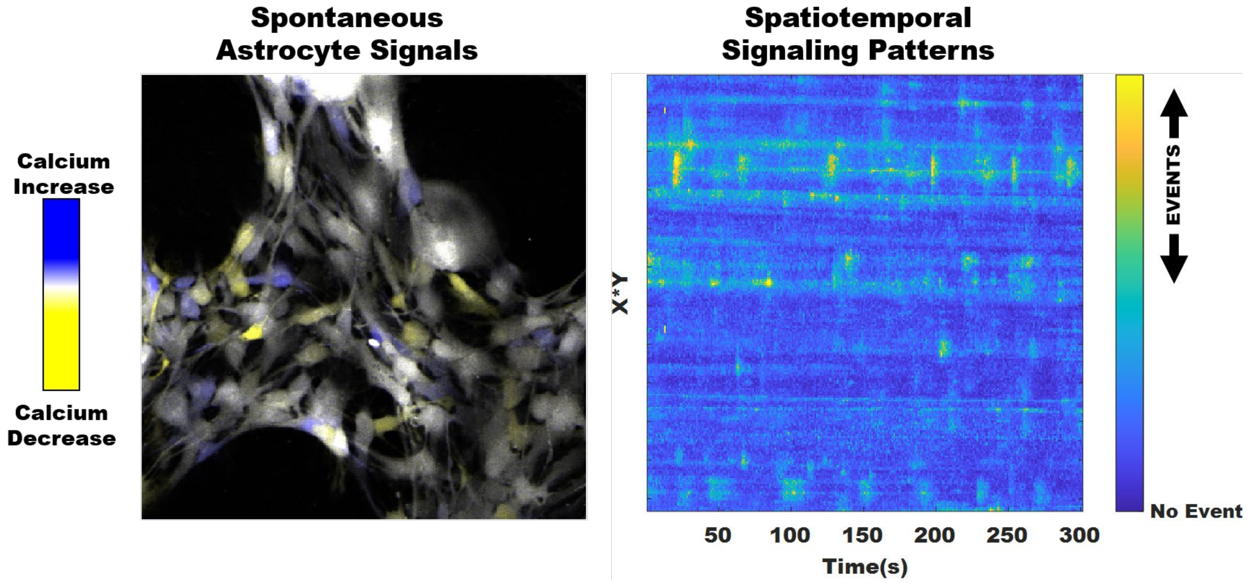


*Fig S1: Spontaneous spatiotemporal calcium patterns. We refigure the snapshots presented in Fig. 1 with a transformed view of the spatiotemporal patterns in astrocyte networks. The spatiotemporal patterns are produced from permuting the dimensions of the data to show time on the x-axis. All calcium events are shown. They are localized in elliptical regions.*

This study does not regard the calcium waves present in astrocyte networks as adaptive mechanisms. Indeed, all astrocyte subtypes in the study spontaneously experience calcium events throughout the network. These complex patterns are shown in a permuted data dimension. This dimension demonstrates the complexity in terms of location, shape, and duration of spontaneous calcium events in astrocyte networks. Information theoretic tools are well-suited to handle the rich, non-linearity present in this data (Fig S1).

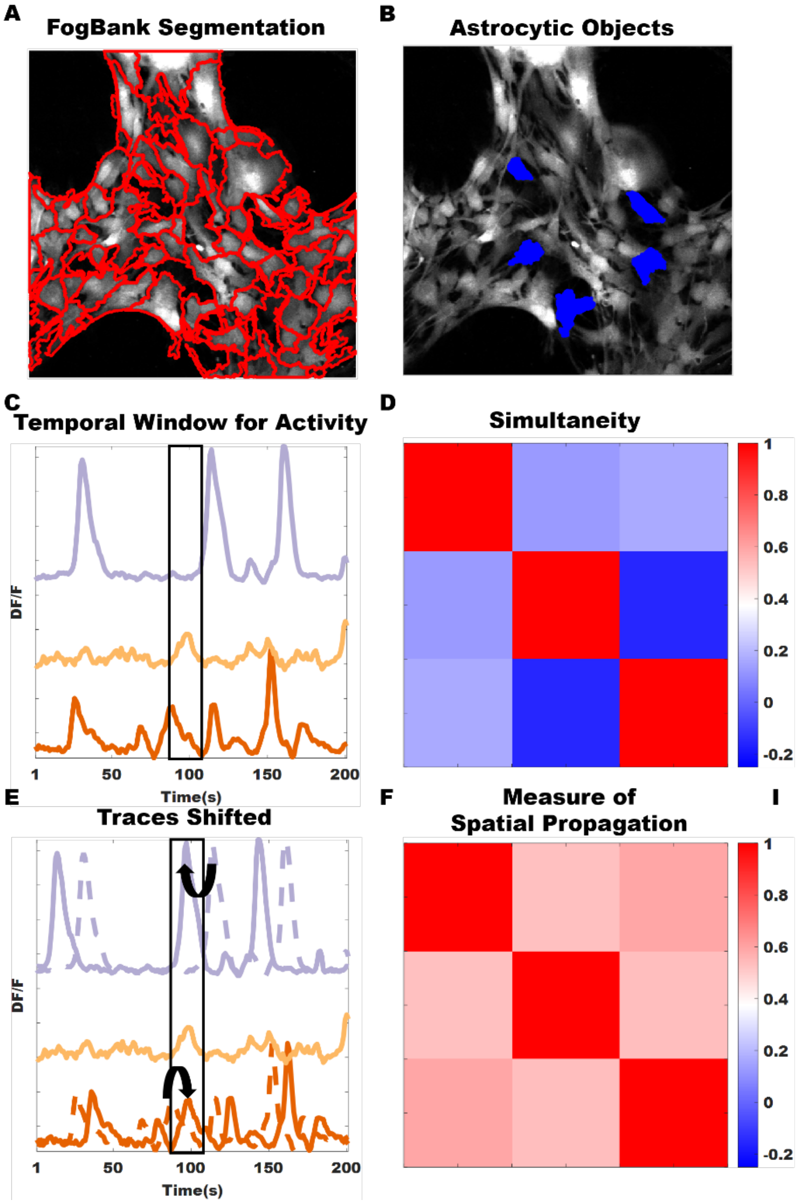


*Fig S2: Construction of objects, and time-delayed correlation versus Pearson. (A) FogBank image shows the boundaries of all segmented cells. For (B) which is reproduced from Fig 2, we show how we went from FogBank to the traces shown in the paper. (C) Demonstrates that for slow traces like astrocytes, Pearson correlation is ineffective in understanding how information is transported. (D) The simultaneity measure of Pearson correlation results in low correlation values. (E) Cross correlation allows traces to be shifted, which makes sense for the slow calcium signals in astrocyte networks. (F) Contrasted to (D), cross correlation values better reflect area overlap in calcium events.*

Astrocytes are segmented using FogBank, a morphological watershed segmentation algorithm[1]. A complete FogBank segmentation is shown in Fig S2A. The examples used in Fig 2 are replicated here in Fig S2B, showing the visual transition from FogBank to the figure as displayed in the text. FogBank segmentation provides distances to each other segmented object (astrocyte); we take the center of mass of each object to find distances between FogBank segmented objects.

The use of correlation for astrocytes is modified from the conventional use of Pearson correlation for neurons. Considering the framerates and the millisecond (ms) timescale for electrical neuronal firing, the use of Pearson correlation (referred in this Supplement as simultaneous firing) is appropriate. The simultaneous firing of neurons allows for Pearson correlation to assess connectivity and other related measures accurately. However, in the case of astrocytes with slow firing on the second timescale, Pearson correlation is inappropriate. Notice in the ‘temporal window for simultaneous firing,’ marked by the bounded rectangle in Fig S2C. Visually, we note that the peaks of calcium events are not aligned which leads to poor correlation output in Fig S2D. However, using Cross Correlation (used in our main text and referred to in this Supplement as a measure of spatial propagation) is appropriate to address the time lags in between calcium events for astrocytes. Shifting the purple (top) and orange (bottom) traces in Fig S2C, shifted traces shown in Fig S2E, we see that the calcium events are more in line, in this case indicating that information was transmitted forward to the yellow (middle) trace; backward information was transmitted from the purple trace to the yellow trace (which equally indicates forward from yellow to purple). Time lags assume positive or negative values, but are taken as positive in the calculation of speed. See improved correlation results, relative to Pearson output, for astrocyte calcium events in Fig S2F.

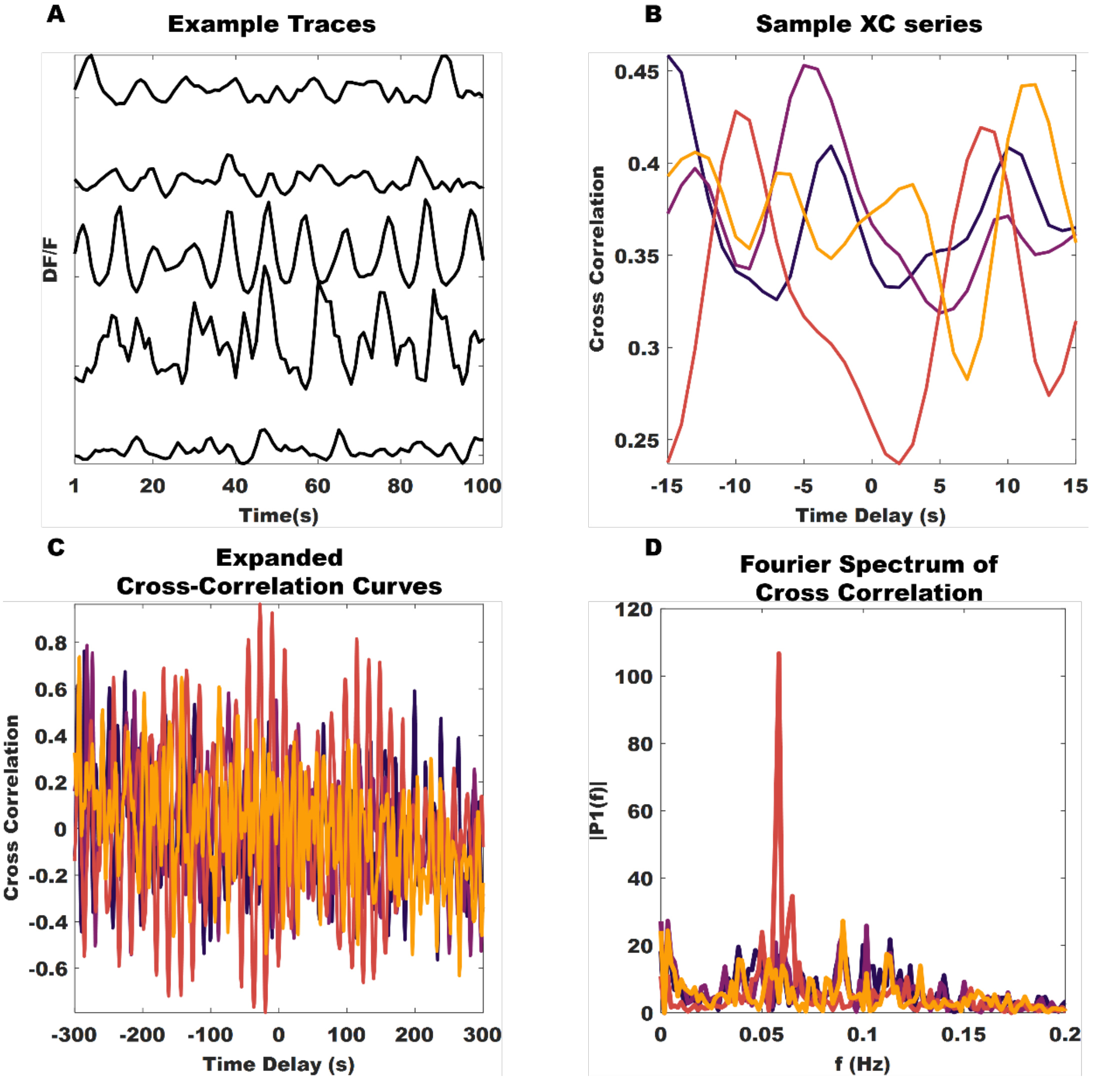


*Fig S3: Analysis of peaks in cross-correlation curves. (A) We reproduced arbitrary stellate time series. (B) For these curves, we report 4 cross-correlation curves associated with time-lags corresponding to the window size used in this study. (C) Expanding*

*this correlation curve to un-physiological time lags, we observe the presence of two modes in this data. (D) Removing the larger mode (subtracting the mean), we find that these peaks associated with maximal correlation are well defined.*

In order to verify that the time lags from either cross-correlation or mutual information are properly extracted, we looked at cross correlations for a wide range of time lags. Using cross-correlation as a representative example, we analyze an arbitrary group of stellate calcium traces in Fig S3A. Some of the cross-correlation curves between these traces are shown in Fig S3B. The curves are smooth, indicating that a well-defined magnitude and time-lag with the largest cross correlation exist. We note that for some curves additional peaks of comparable height are seen. As shown in Figs S3C and S3D, these additional peaks are consistent with the period character of many calcium traces, which yields well-defined peaks in the Fourier spectrum (Fig S3D). Corresponding to the peak in Fig S3D at 0.05Hz (indicating a 20 second periodicity), we find that the corresponding cross correlation (red curve in Fig S3B) has two peaks 20 seconds apart. We systematically choose the largest peak to reflect that most likely time delay and magnitude of cross-correlation.

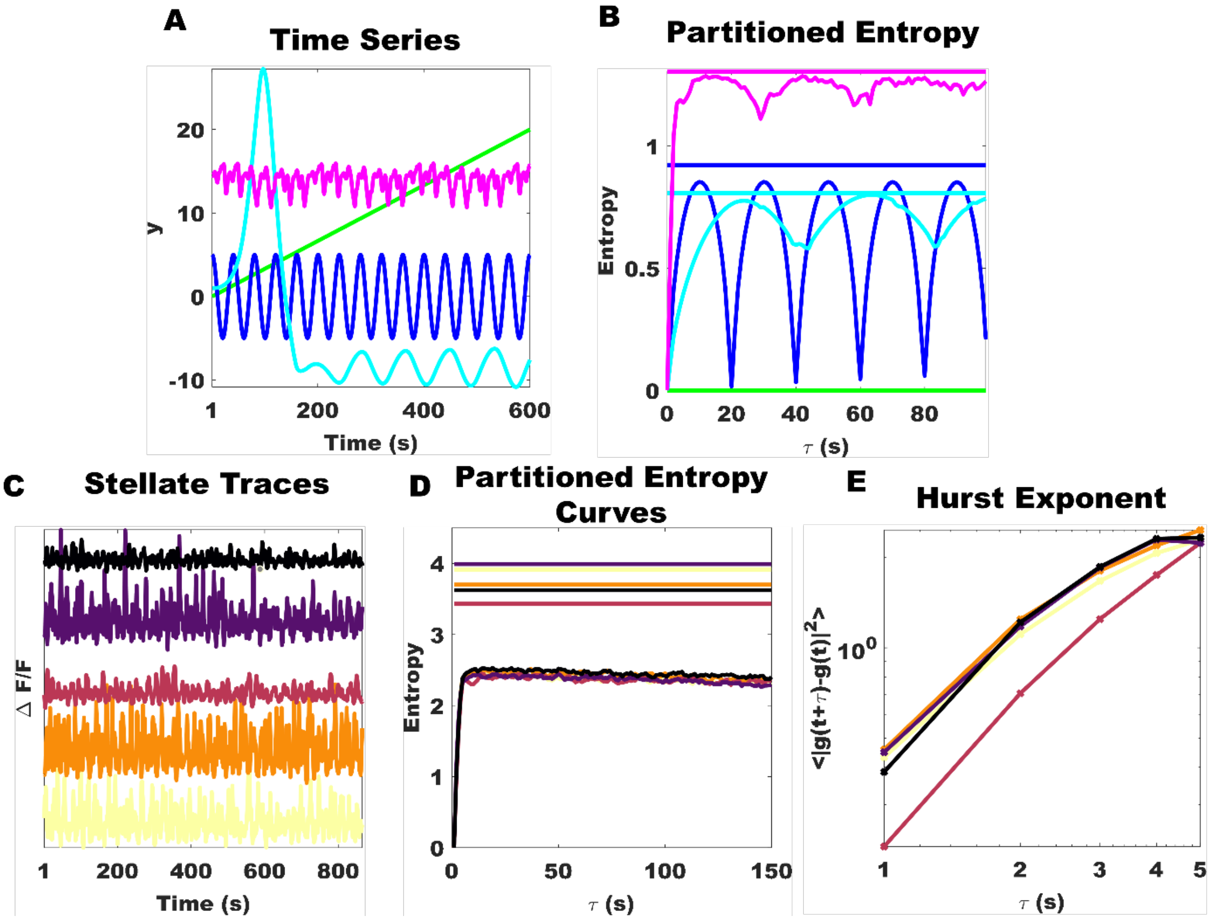


*Fig S4: We reproduced a partitioned entropy figure to demonstrate our code is sound. We plot representative curves for partitioned entropy. Since the partitioned entropy ratio retains the same bounds ([0 1]) are the Hurst exponent, for completeness we show how these are difference metrics. Hurst exponent is finding the slope of the graph shown, loglog plotted, in (E). τ shown in (D) refers to symbol state time whereas in (E) it reflects acquisition rate of the image sequences.*

Partitioned entropy enables a more individualistic interpretation of the calcium events of astrocytes. In order to correctly ensure the measure, here used as a comparative metric to assess difference between astrocyte subtypes, is used appropriately we replicate the results in Shiozawa et al.[2] In Fig S4A, for a straight line (green), a sine curve (blue), the Lorenz equation (cyan), and Thomas' cyclically symmetric attractor (pink), we replicate partitioned entropy curves. The maximal entropies for these curves are indicated by the zero slope lines with the same color as the partitioned entropy curve in Fig S4B. As one can see, there is no partitioned entropy for a line, the partitioned entropy for a sine curve is roughly 0.5 (as indicated by the sinusoidal nature of the partitioned entropy curve), whereas for the Lorenz and Thomas systems, the partitioned entropy curves saturate closer to maximal entropy values, indicating that these time series are chaotic.

For this paper, we reproduce some stellate traces (Fig S4C) with respective partitioned entropy curves (Fig S4D). We note that these curves visually appear comparable to chaotic partitioned entropy curves; however, these curves hover around ½ the maximal entropy value. Exploring the nature of chaos in astrocyte systems is beyond the scope of this paper, but worthy of further investigation.

Since the partitioned entropy ratio and the Hurst exponent assume comparable values, [0 1], we reproduce the curves for evaluating the Hurst exponent for the stellate traces in Fig S4C. As shown in Fig S4E, the Hurst exponent is the slope of the log-log plot between the expected squared difference between values in a time series offset by $\tau$ and $\tau$ itself. There is a connection between the Hurst exponent and partitioned entropy, but these metrics are distinct and the values of 0.5 signify different physical meaning.

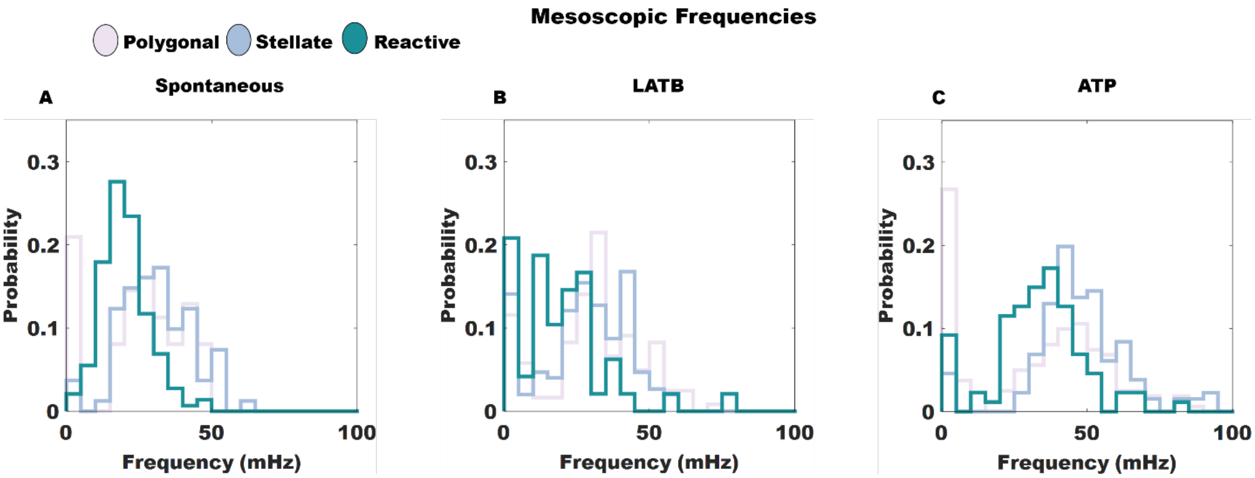


*Fig S5: Fourier frequencies. Peak frequencies for individual traces for polygonal, stellate, and reactive traces in (A) spontaneous conditions, (B) LATB conditions, and (C) ATP conditions are shown for completeness.*

Due to the nonlinearity of astrocyte signals and the use of symbolization, finding the frequencies of these traces does not logically flow with the structure of the main text. The nature of astrocyte calcium events renders any output frequency a dubious value, i.e. astrocytes need not, and do not, obey the same frequency within certain distinct time windows in a series. However, for completeness, we report these values. The values reported are the peak frequency associated with the periodogram of individual traces. Nonetheless, we find that reactive astrocytes have the lowest spontaneous frequency with stellate having the highest (Fig S5A). Latrunculin B increases polygonal and stellate distributions while lowering reactive ones (Fig S5B). ATP increases the frequencies of all astrocyte subtypes (Fig S5C).

1. Chalfoun, J. *et al.* FogBank: a single cell segmentation across multiple cell lines and image modalities. *BMC Bioinformatics* **15**, 431 (2014).

2. Shiozawa, K., Uemura, T. & Tokuda, I. T. Detecting the dynamical instability of complex time series via partitioned entropy. *Phys. Rev. E* **107**, 014207 (2023).
